# Morph-linked variation in female pheromone signaling and male response in a polymorphic moth

**DOI:** 10.1101/2024.07.13.603243

**Authors:** Chiara De Pasqual, Eetu Selenius, Emily Burdfield-Steel, Johanna Mappes

**Affiliations:** Organismal and Evolutionary Biology Research Program, University of Helsinki, 00014 Helsinki, Finland; Institute of Biodiversity and Ecosystem Dynamics (IBED), University of Amsterdam, Amsterdam 1098 XH, The Netherlands

**Keywords:** color polymorphism, pheromone variation, receiver variation, sexual selection, wood tiger moth

## Abstract

1. Understanding the maintenance of genetic variation in reproductive strategies and polymorphisms in the wild requires a comprehensive examination of the complex interactions between genetic basis, behavior, and environmental factors.
2. We tested the association between three color genotypes and variation in female pheromone signaling and male antennal morphology in the wood tiger moth (*Arctia plantaginis*). These moths have genetically determined white (WW, Wy) and yellow (yy) hindwings that are linked to mating success and fitness, with heterozygotes (Wy) having an advantage. We hypothesized that attractiveness and reproductive success are correlated, with Wy females being more attractive than the other two genotypes which could contribute to maintaining the polymorphism.
3. Female attractiveness was tested by baiting traps with females of the three color genotypes both in low-(i.e., field setup) and in high-population density (i.e., large enclosure setup). Male’s ability to reach females was correlated to their own color genotype and antennal morphology (length, area, and lamellae count).
4. Contrary to our prediction, morph-related reproductive success and attractiveness were not correlated. Heavier Wy females attracted a lower proportion of males compared to WW and yy females. Specifically, an increase in weight corresponded to a decreased Wy but increased yy female attractiveness. yy females were generally more attractive than others likely due to earlier pheromone release. In males, lamellae count and genetic color morph were linked to the male’s ability to locate females. Furthermore, male traits affected their ability to reach females in a context-specific way. Males with denser antennae (i.e., higher lamellae count) and white males reached the females faster than yellows in the enclosure, while yellow males located females faster than whites in the field.
5. Our results indicate that higher yy female attractiveness was likely affected by the combined effect of early pheromone release, female weight and higher population density. Males’ searching success was affected by morph-specific behavioral strategies and local population density. Ultimately, the combined effect of genotype-related pheromone signaling strategies of females together with environment-dependent male behavior affect male response and potentially contribute to maintaining variation in fitness-related traits.

## Introduction

Phenotypic variation is often maintained by intricate interactions among multiple selective pressures. Genetic correlation between different sets of traits can significantly influence individual fitness and performance, thereby maintaining phenotypic variability (McKinnon & Pierotti, 2010). For example, in lizards, correlations between color morphs and chemical signals have been observed to facilitate assortative pairing (Pellitteri-Rosa et al., 2014). Similarly, disassortative mating between white and tan morphs in the white-throated sparrow (*Zonotrichia albicollis*) preserves both plumage colorations in populations (Tuttle et al., 2016). Such trait variability, especially in sexual signals and secondary sexual traits, serves as the foundation for mate choice processes (Andersson, 1994; Fisher, 1930; Johansson & Jones, 2007). While extensive research has explored how variation in sexual traits of signalers (e.g., female sex pheromones) influence mate choice (e.g., De Pasqual et al., 2021; Johansson & Jones, 2007; Steiger & Stökl, 2014; Groot et al., 2014), there has been relatively limited investigation into how variation in the sensory structures of receivers (e.g., male antennae) contribute to mate choice (Elgar et al., 2019; Jayaweera & Barry, 2017). Given that both sexes interact in the wild, sexual selection likely results from the choices made by both sexes. However, how variation in signaler and receiver traits, and in the consequent mate responses and choices, relate to particular genotypes remains relatively unknown. In this study, we address this overarching question by investigating how the interplay between the morph of the signaler and the variation in receiver’s sexual traits can influence mate responses and choices, ultimately contributing to the maintenance of genetic diversity in reproductive traits and polymorphisms under precopulatory selection.

In moths, females typically signal to potential mates by releasing sex pheromone. Pheromones can be costly to produce and maintain (Blankers et al., 2021; Harari et al., 2011) and variation occurs both within the individual lifespan and between individuals (De Pasqual et al., 2021; Blankers et al., 2021; Pham et al., 2021). Through pheromone variation and the use of signaling strategies, females can exert sexual selection on males by manipulating the arrival of males (Greenfield, 1981; Johnson et al., 2017a) and alter intrasexual dynamics by outcompeting same-sex conspecifics (Pham et al., 2020, 2021). Females can temporally change their signaling strategy by increasing their calling (i.e., when a female releases sex pheromone) effort with age to reduce the chances of dying unmated (Greenfield, 1981; Johnson et al., 2017a; Umbers et al., 2015) or adjust their signaling effort in accordance with the perceived level of intrasexual competition. For instance, *Uraba lugens* females that develop in high-density larval environments call earlier and for longer compared to females that develop in low-density environments (Pham et al., 2020).

While there is mounting evidence of the ecological importance of pheromone variation and signaling strategies for intrapopulation dynamics (De Pasqual et al., 2021; Johansson & Jones, 2007; Steiger & Stökl, 2014), the effect of natural variation of sensory structures (i.e., antennae) on signal detection, receiver behavior, and its effects on scramble competition, is seldom investigated (Elgar et al., 2019; Jayaweera & Barry, 2017). The minute quantities of sex pheromone released by females (Symonds & Elgar, 2008; Wyatt, 2014) exert selection on male antenna for faster mate location, which is generally achieved through bigger antennae (Elgar et al., 2019; Johnson et al., 2017a; Jayaweera & Barry, 2017; Symonds et al., 2012). Antennal development is, however, costly for males (Johnson et al., 2017b; Pham et al., 2022; Symonds et al., 2012), thus antenna morphological variation is likely to affect male’s ability to locate females and mating success. For instance, males of the diamondback moth (*Plutella xylostella*) with partially or completely ablated antenna experienced a steep decrease in their mating rate (Yan et al., 2014) and males of the false garden mantid (*Pseudomantis albofimbriata*) took longer to locate females the lower the number of sensilla on their antennae (Jayaweera & Barry, 2017). While knowledge on the ecological effect of variation both in the signaler and receiver’s trait is increasingly accumulating (De Pasqual et al., 2021; Jayaweera & Barry, 2017; Johansson & Jones, 2007; Johnson et al., 2017a; Pham et al., 2021; Umbers et al., 2015) there is yet very little understanding on how such variation can interact to affect the mate choice processes and maintain phenotypic trait variation in populations.

Due to its polymorphic hindwing coloration and mate recruitment based on a long-range sex pheromone, the wood tiger moth (*Arctia plantaginis*) is a promising study organism to investigate the relative contribution of females and males’ signaler and receiver trait variation during mate recruitment (i.e., precopulatory stage). Male hindwing coloration is genetically determined by a one locus-two allele polymorphism (Brien et al., 2022; Nokelainen et al., 2022; Suomalainen, 1938). The dominant white (W) and recessive yellow (y) alleles determine three color genotypes (WW, Wy, yy) which are phenotypically expressed as white or yellow discrete hindwing coloration. Females carry the male color alleles but do not phenotypically express them as their hindwing coloration varies continuously from yellow to red (Lindstedt et al., 2011; Nokelainen et al., 2022). Previous studies investigating morph-specific associations at the copulatory and postcopulatory stage found that morph- and genotype-linked advantages are context-dependent rather than fixed across ecological contexts. Females generally prefer to mate with white males (Gordon et al., 2018; Nokelainen et al., 2012) but their preference seems to be flexible and affected by the male morph frequency. Males of either morph have a reproductive advantage when they represent the most common morph (Gordon et al., 2015) whereas all male morphs have equal chances of mating when females are not presented with an alternative mating partner (Chargé et al., 2016; De Pasqual et al., 2022). Yellow males are generally characterized by lower reproductive success than whites (De Pasqual et al., 2022; Gordon et al., 2018) while Wy females have the highest reproductive output among female genotypes (De Pasqual et al., 2022). As a proxy for precopulatory mate searching behavior, Rojas et al., (2015) found that yellow males have a narrower daily flying activity than white males, whereas very little is known about how female behavior and attractiveness vary.

In this study, we explore the links between color-related genotypes, signaling behavior, receiver morphology and mating responses in the context of precopulatory mate choice. In particular, we studied how signalers (i.e., females) differed in their attractiveness to males and how receivers (i.e., males) differed in their ability to reach the females. First, we tested whether females of the three color genotypes were differentially attractive to males. We hypothesized that heterozygote females with highest reproductive success are most attractive, following De Pasqual et al., (2022). Since females were indeed differentially attractive, we tested for potential signaling strategies, by investigating how long it took for the female to attract the first male and the duration of female attractiveness. Second, we tested whether male color genotype or antennal morphology correlated with their ability to locate a female. We hypothesized that white males would reach females faster than yellow males, based on Rojas et al., (2015). Finally, we tested the hypothesis that females attracted males bearing different color alleles from their own, indicating that disassortative attractiveness could contribute to maintaining both color alleles (W, y). These experiments were carried out both in the field and semi-field setup (i.e., large net-covered enclosure), to mimic a scenario of low and high density population, respectively.

## Materials and methods

### Study species

The lab-reared moths used in these experiments came from the laboratory stock established in 2013 from Finnish wild caught individuals and maintained at the Department of Biological and Environmental Sciences at the University of Jyväskylä (Finland). Three genotype lines (i.e., WW, Wy, yy) have been established for experimental purposes since the moths’ color genotypes cannot be inferred reliably by simply looking at their hindwing coloration (Nokelainen et al., 2022). The genotype lines are maintained through controlled matings that aim to keep genetic variability as high as possible. This is achieved by forming mating pairs of all possible color genotype combinations selecting genotyped individuals from as many families as are available in the stock. The moths are reared by following the general husbandry protocol described in the Supporting Information of this paper, and previously reported in Nokelainen et al., (2022) and De Pasqual et al., (2022). After eclosion, adult moths were placed in a cooler cabinet (7°C) and were taken out around 12.00 p.m. to allow the individual to acclimatize at room temperature before being tested.

Females signal to potential males by releasing sex pheromone (a behavior commonly referred to as ‘calling’) from a gland located at the tip of their abdomen. Once males detect the female sex pheromone, they fly upwind to reach the pheromone source. Under laboratory conditions, females release pheromone daily for about four hours (mean 228 min ± 122 SD) starting ca. 1.5 hours before the sunset (on the day after their eclosion). Males actively search for potential mating partners between 5.00 p.m. and 10.00 p.m. (Gordon et al., 2015; Nokelainen et al., 2012) with mating activity peaking between 10.00 p.m. and 11.00 p.m. under laboratory conditions. The wood tiger moth is a capital breeding species; adults do not feed, therefore larval diet is particularly important both for their development and for the adult stage (e.g., egg numbers, sperm quality, sex pheromones) (Tammaru & Haukioja, 1996).

### General methods

Potential associations between genotype-related variation in female pheromone signaling and male antennal morphology were tested in two experimental setups. In both setups, females were placed in bucket traps (Pherobank cat. No. 30202) from where they could signal potential males through the release of sex pheromone, but the female could not be seen and physical contact was prevented by a fine mesh covering the housing cage. Females were tested over one night and replaced with new females for the following night. Males that were attracted would fall into the trap and be collected on fly strips (Raid, product no. 100030684) at the bottom of the trap-bucket.

#### a) Field setup

Traps were set in six locations within Tornio municipality (65.8473°N, 24.1520°E), Northern Finland. These locations were chosen based on previous *A. plantaginis* sightings registered in the Finnish Biodiversity Information Facility (laji.fi) (Fig. 1). In each location, we tested two lab-females of different genotypes placed in two separate traps per location and in the following genotype combinations: WW & Wy, WW & yy, Wy & yy (Fig. 1). We tested these three combinations of genotypes 13, 12 and 9 times respectively, and retained 11, 11, and 9 trials, for each combination, respectively. In the excluded trials, one of the two females was found dead the following morning, and we could not exclude the possibility that the lack of attractiveness was simply due to her death. Females did not significantly differ in weight (F_(2,58)_ = 0.02, *P* = 0.98) or age (F_(2,59)_ = 0.34, *P* = 0.71) between genotypes.

**FIGURE 1.**
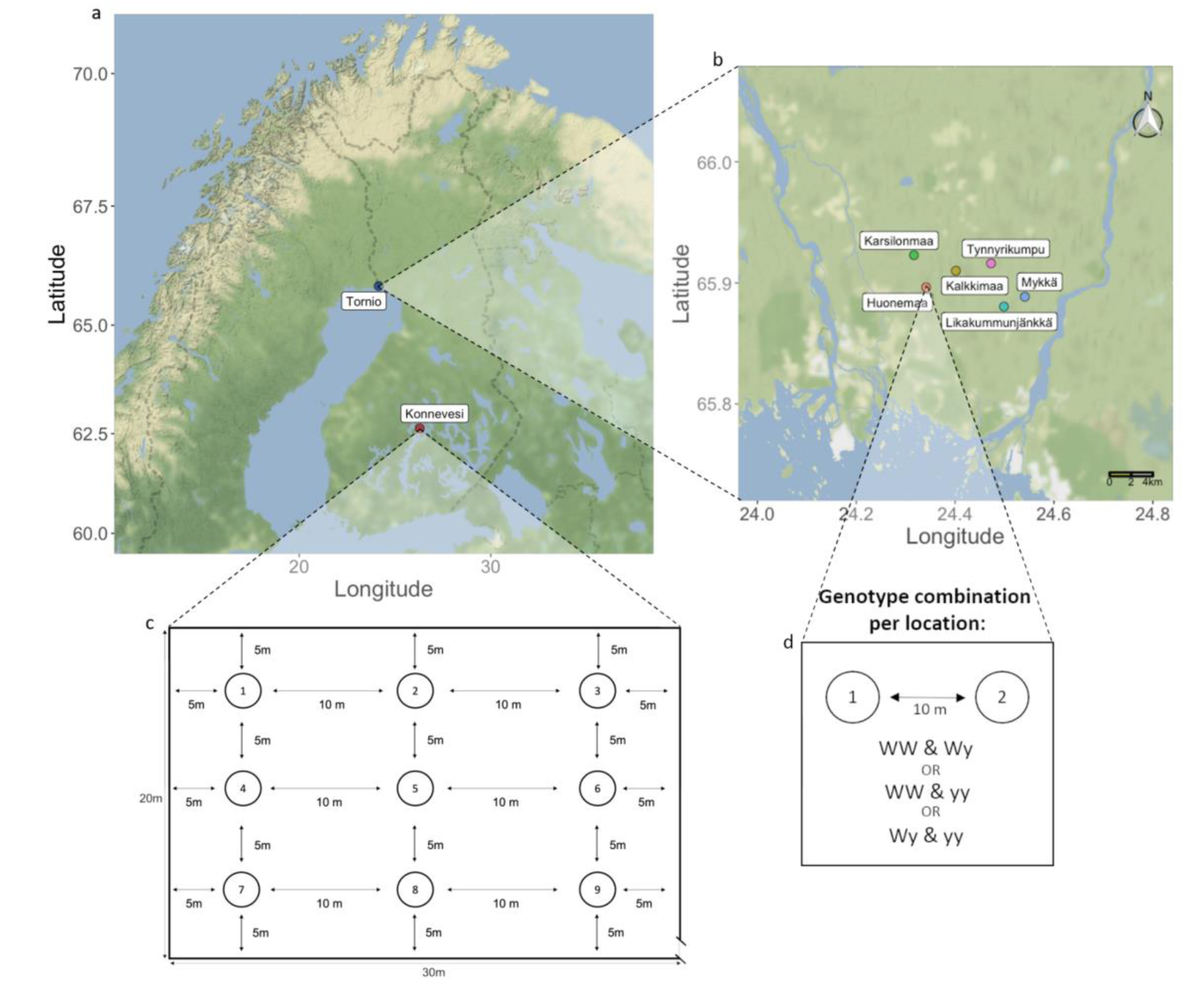
a) The map shows the locations of the two experiments. Female attractiveness and male’s ability to locate females were tested b) in a field setup by deploying traps in six locations in Tornio municipality and c) in an outdoor enclosure experiment in the Konnevesi Research Station. d) shows the trap position and genotype combinations per location in the field setup. The numbers in circles denote the trap position in c) the enclosure and d) the field setup.

Each genotype combination was randomly assigned to a location on the first day, after which they were rotated between locations, to ensure that each location would house each pair combination at least once. The traps were placed approximately 10 m apart and hung at about 90 cm from the ground, and genotypes were rotated between the ‘right’ and ‘left’ position to avoid position biases. Traps were set between 10.00 a.m. and 2:00 p.m. and female attractiveness was tested overnight. The following day, new females were placed in the traps after removing the old ones and the wild-caught males found in the traps were collected.

#### b) Enclosure

Female attractiveness was also tested in a 30 x 20 x 3 m enclosure at the Konnevesi Research Station (62.6278°N, 26.2939°E, Jyväskylä municipality, Central Finland) with lab-reared males and females. Traps were placed in a 3 x 3 grid on 90 cm high poles which were placed at five to 10 m distance (Fig. 1). Males were individually marked with a four-dot two-color system that allowed the identification of individual males caught in the trap. Females were introduced into the enclosure at 3.30 p.m. and males were released at 4:00 p.m. from four releasing points to avoid position biases.

Twenty males per genotype (n = 60) were released every second night, while a new set of three females per genotype (n = 9) was tested every night (except for the first two female replicates, in which the same set of females was tested over two nights). Female genotype position was rotated at every replicate to avoid position biases. A total of 180 males were released over six nights and a total of 54 females (with 45 unique females) were tested for their attractiveness. Male weight (F_(2,182)_ = 0.67, *P* = 0.51), male age (F_(2,189)_ = 0.52, *P* = 0.59) and female age (F_(2,51)_ = 1.38, *P* = 0.26) did not differ between genotypes. Female weight significantly differed between genotypes (F_(2,49)_ = 6.93, *P* = 0.002) with yy females being significantly heavier than WW (TukeyHSD; CI (8.27, 50.63), *P* = 0.004) and Wy females (TukeyHSD; CI (5.57, 49.27), *P* = 0.01). For this reason and due to collinearity between weight and genotype (VIF = 7.26), we normalized the variable weight by subtracting the mean weight per genotype and dividing by the standard deviation of all female weight (Fig. S1). A few males were already present in the enclosure from a previous experiment. Among the 36 males caught from the previous experiment, we collected four WW, three Wy, and five yy, while the remaining 23 could not be genotyped, but were considered for the total individual female attractiveness. Traps were checked at two-hour intervals, at 6.00 p.m., 8.00 p.m., 10.00 p.m., and 12.00 a.m., at which point the moths were no longer active. At each time-interval, males that were found in traps were collected.

### Female attractiveness

Potential associations between female color genotype and sex pheromone attractiveness were tested considering the likelihood of attracting at least one male and the overall attractiveness of the female color genotypes (i.e., the number and the proportion of males caught). In addition, we compared the daily proportion of males attracted by female genotype in both setups. In the enclosure, we tested how quickly a female genotype would attract males within a night, and whether the proportion of males attracted was affected by the duration of female attraction. Finally, we tested for any sign of disassortative attractiveness by considering both the male genotype and/or color phenotype attracted by each female genotype.

### Male order of arrival

In both the field and the enclosure, we tested whether male color genotype and antennal morphology correlated with their order of arrival to the traps by considering their genotype/ color morph and antenna morphology. For the males caught in the field we recorded their hindwing color morph (i.e., white or yellow), and by reading the four-dot two-color marking for the males collected in the enclosure, we had information about their color genotype, weight, and age. From the males that presented at least one fully intact antenna, we excised one antenna as close as possible to the head and stored it in ethanol (80%). Eighteen males collected from the field and 31 males tested in the enclosure lacked any intact antennae, therefore were excluded from the analyses. Each antenna was then flattened onto a glass slide covered with millimeter paper and photographed under a microscope (1 x magnification, camera 14.0MP Japan CMOS and software S-EYE) (Fig. S2). The area and length of the antenna were measured using ImageJ software. We first set the scale by measuring the millimeter paper. We then measured the area by including the flagellum and one side of the lamellae. The length was measured as the length of the flagellum. The number of lamellae was manually counted (Fig. S3). In total we measured 139 wild- and 122 lab-reared male antennae.

### Statistical analyses

All analyses were performed using R (v.4.1.2) and RStudio (v.2022.02.3). In all cases where we fit two models, one with an interactive effect between two factors and one without, the final model was selected based on the lowest AIC. All the fit models (with and without interactions) and the models selected based on lowest AIC are reported in Table S1. Once the final model was selected, the overall effect of the fixed factors was tested with Chi-square implemented with the ‘anova’ function (‘car’ package, v. 3.0-12).

### Female attractiveness

#### a) Field setup

To test for the effect of female genotype, night of experiment and weight, on the likelihood of attracting at least one male we fit two Generalized Linear Mixed Models (GLMM; ‘glmer’ function, ‘lme4’ package v.1.1-27.1) modelled with a binomial distribution, one including the interaction “female genotype*night of experiment”, and one without the interaction (Table S1a). To test for the effect of female genotype, night of experiment, and weight (fixed factors) on the number of males attracted (response variable), we fit a GLMM model, with the response variable “male count” modelled with Poisson distribution (Table S1a). Because of the pairwise design, we created an identifier that was specific to each trial by assigning to each female a label made of the night and location of testing. This allowed us to correct for the frequencies of males that were attracted by two female genotypes, as each night and location included in the identifier corresponded to one pair of female genotypes. This label was then set as a random effect. The night variable in the interaction term was treated as discrete factor rather than continuous covariate since we were interested in testing all the possible night and genotype combinations. In addition, we tested for different attractiveness of the female genotypes by testing the daily number of males attracted by female genotype by setting the interaction between female genotype and night (fixed factor) in a GLM with Poisson distribution. The total number of males caught per night was used as a covariate. This analysis was performed with the genotype combinations of night 2, 3 and 4 as these nights had an equal number of females tested (unlike night one where some females were excluded as they were found dead the day after). The night variable was treated as a discrete factor as before. Finally, we tested whether females attracted males bearing different color alleles from their own (i.e., disassortative attractiveness). For each female, we considered the number of white and yellow males that she attracted. Thus, we tested for the interactive effect of female genotype and male morph (fixed effect) on the number of males attracted by females (response variable) and controlled for the effect of night, location and female ID (random effects) by fitting a GLMM, modelled with Poisson distribution.

#### b) Enclosure

We tested whether the likelihood of attracting at least one male (response variable 1) and the proportion of males caught (response variable 2) were affected by female genotype and (normalized) weight (fixed factors), and we controlled for the trap position and replicate (random factors) by fitting four GLMM models, two per response variable, with binomial distributions. One model per response variable included the interaction between “female genotype*weight” (Table S1b). The proportion of males caught was calculated as the number of males attracted by the female over the number of males present in the tent at the start of the replicate through the function ‘cbind’. In addition, as for the field setup, we tested for different attractiveness of the female genotypes by testing the daily number of males attracted by female genotype by setting the interaction between female genotype and replicate (fixed factor) in a GLM with Poisson distribution. The total number of males caught per replicate was used as a covariate. The variable replicate in the interaction term was treated as a discrete factor rather than continuous covariate since we were interested in testing all the possible replicate and genotype combinations. Finally, we tested for disassortative attractiveness by using a similar approach to the field setup. For each female, we considered the number of WW, Wy and yy males that she attracted. Thus, we tested for the interactive effect of both male and female genotype (fixed factor) on the proportion of males attracted (response variable), by fitting a GLMM model with the response variable modelled with binomial distribution and controlled for the effect of replicate and female ID (random effects).

##### Strategies of female attractiveness

Since females were differently attractive in the enclosure setup, and the same female was observed across multiple time-check points throughout the night, we tested for differences in signalling strategies associated with time of onset and duration of calling. First, we tested whether female genotype (fixed factor) affected how long it took for each female to attract their first male (for simplicity called ‘onset’) by fitting a Cox Proportional Hazard Model (henceforth ‘Cox model’) (‘coxph’ function, ‘survival’ package v. 3.2-11). At each time-checkpoint (i.e., 6.00 p.m., 8.00 p.m., 10.00 p.m., 12.00 a.m.) traps were checked for males and emptied. If a male was found in the trap during the first time-checkpoint, we assigned value 1 to the female. If the first male in the trap was found in the second time-checkpoint, the female would be assigned value 2, and so on until the last time-checkpoint of the evening (max value assigned was 4). Finally, we tested if earlier onset (a proxy for calling -denoted as 1 to 4 as for the previous analysis) or longer duration of female activity (a proxy for duration - calculated as the total number of time-checkpoints males were found in the trap as there were no instances in which males would be found in a female’s trap on discontinuous time-checkpoints) (fixed factors) affected the number of males attracted (response variable). We thus fit two GLMMs with Poisson distribution, one with the interaction ‘onset * duration of female activity’ and one without. The number of males in the tent was set as covariate and we controlled for the effect of replicate and trap position (random effects) (Table S1c).

### Male order of arrival

#### Antennal morphology

Before delving into the potential role of male trait variation in affecting the order of arrival to traps, we checked for potential differences in antennae due to color genotype/morph, weight and male origin (i.e., wild-caught or lab-reared). In addition, we tested for collinearity between antennal length, area and number of lamellae. The analyses are described in the Supporting Information.

#### Effect of antennal morphology and genetic color morph on the order of arrival

To test for the effect of antenna morphology (i.e., length, area, number of lamellae), male morph (i.e., white and yellow), and their interactions (fixed factors) on the order of wild male arrival to traps (response variable) we fit five Cox models, testing all antennal characteristics and male morph interactive combinations (see Table S1d). For lab-reared males, we tested the effect of antennal morphology (i.e., length, area, number of lamellae), male genotype (i.e., WW, Wy, yy), all their possible interactions, weight and age (fixed factors) on the order of arrival to traps, by fitting five additional Cox models (see Table S1e). Finally, we tested for differences in the lamellae number (response variable) of the males attracted by females of the three different genotypes (fixed factor). We therefore fit a GLMM, with Poisson distribution.

## Results

### Female attractiveness

#### a) Field setup

The likelihood of a female attracting at least one male was not affected by the genotype (χ^2^ = 0.15, df = 2, *P* = 0.93), night (χ^2^ = 0.59, df = 1, *P* = 0.44) or weight (χ^2^ = 1.02, df = 1, *P* = 0.31) (Table S2a). While genotype (χ^2^ = 3.03, df = 2, *P* = 0.22) and weight (χ^2^ = 3.51, df = 1, *P* = 0.06) did not significantly affect the number of males attracted, the night of experiment did (χ^2^ = 15.33, df = 6, *P* = 0.02). The number of males attracted decreased significantly over the course of the experiment (see Table S2b). When we tested for different daily attractiveness of the female genotype, the number of males attracted per night was not affected either by the interaction between female genotype and night of experiment (χ^2^ = 3.52, df = 4, P = 0.47), or female genotype (χ^2^ = 6.10, df = 2, P = 0.05) or the night of experiment (χ^2^ = 0.46, df = 1, P = 0.50) or the number of males caught by night, suggesting that all genotypes attracted equally throughout the nights (Table S2c). Finally, we did not find evidence of disassortative attractiveness based on the male morph attracted by female genotype (χ^2^ = 3.50, df = 2, *P* = 0.17) (Table S2d and Table S3).

#### b) Enclosure

The likelihood of attracting at least one male was not affected by female genotype (χ^2^ = 0.21, df = 2, *P* = 0.91) or weight (χ^2^ = 0.01, df = 1, *P* = 0.92) (Table S4a). However, the proportion of males caught was affected by the interaction ‘female genotype*weight’ (χ^2^ = 10.94, df = 2, *P* = 0.004) (Fig. 2). Heavier Wy females attracted a significantly lower proportion of males than WW (GLMM; estimate = 0.99 ± 0.47, z = −2.21, *P* = 0.03) and yy (GLMM; estimate = 1.88 ± 0.59, z = −3.20, *P* = 0.001) females (Fig. 2). While weight did not affect WW females which showed an almost constant slope (0.20), weight affected Wy and yy females in opposite ways, with Wy females being negatively affected (slope −0.79) and yy females were positively (slope 1.09) affected by higher weight (Fig. 2, Table 1). The daily number of males attracted by female genotype was affected by the night of experiment (χ^2^ = 24.42, df = 10, P = 0.001) (Fig. 3), suggesting night-specific female advantages. yy females were significantly more attractive than WW and Wy females in the first two replicates, were significantly more attractive than WW females in the third replicate and showed a tendency of higher attractiveness in the third to fifth replicate. This advantage did not hold for the last replicate, where they were equally as attractive as the WW and Wy females (Table 2; Fig. 3). Finally, we did not find any sign of disassortative attractiveness based on the color genotypes (χ^2^ = 2.20, df = 4, *P* = 0.70) (Table S3 and Table S4b).

**FIGURE 2.**
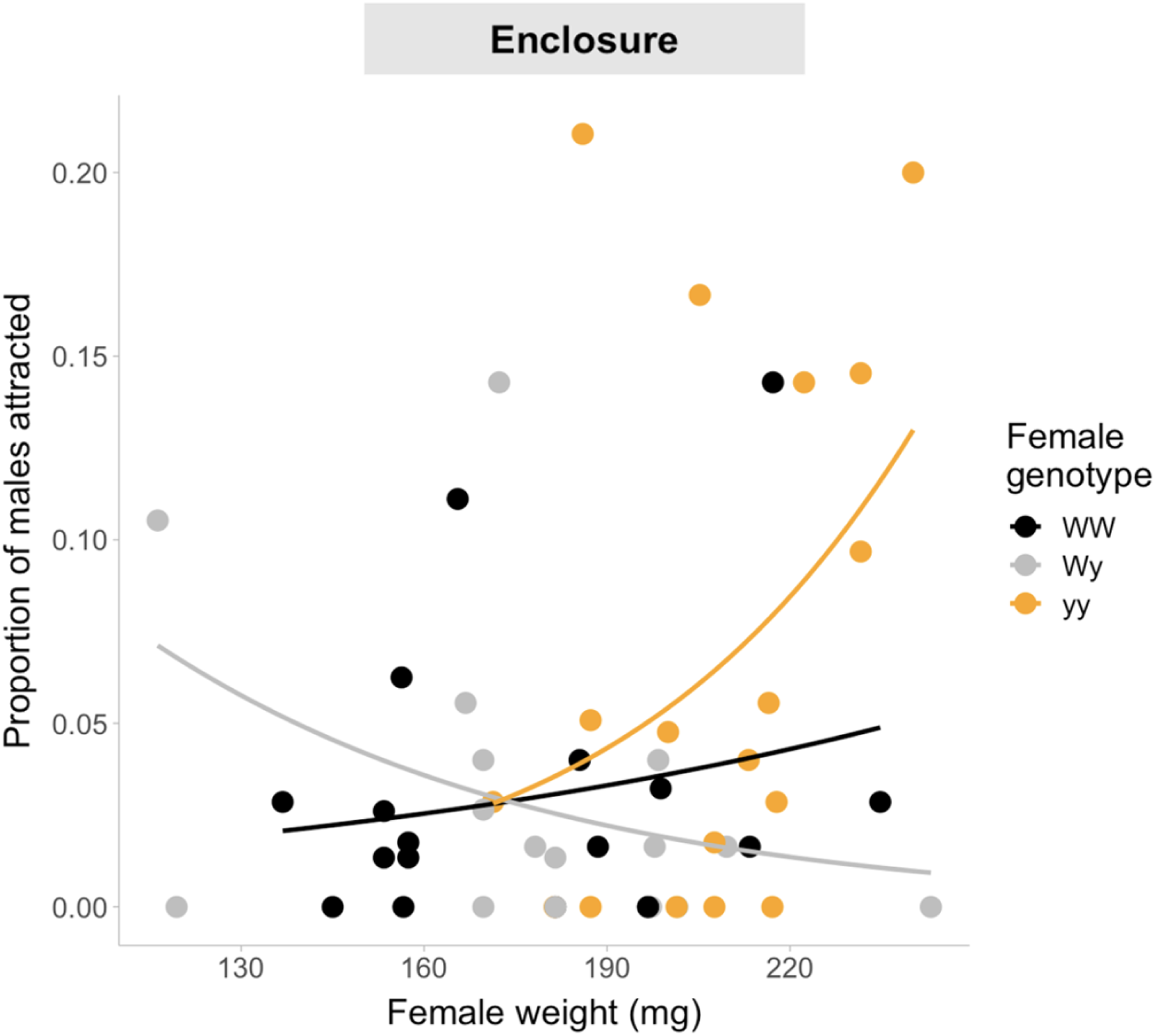
Proportion of males attracted (y-axis) by female genotype in relation to female weight (x-axis) in the enclosure setup.

**FIGURE 3.**
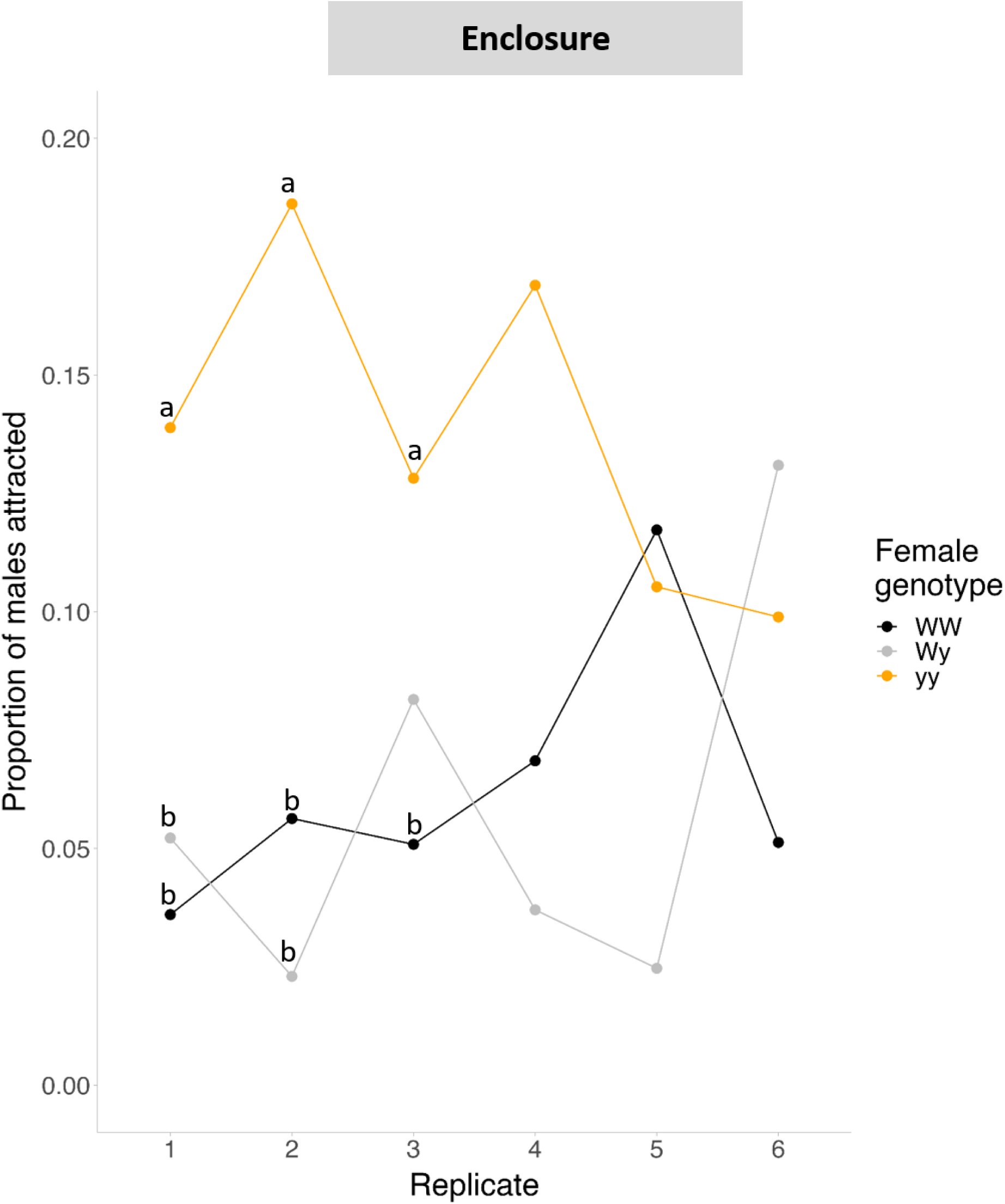
The plot shows the proportion of males attracted by female genotype (y-axis) across the male replicates (x-axis). The significant differences are based on pairwise comparisons on estimated marginal means (EMMs). Significant differences (P < 0.05) by night between genotypes are denoted with letters and the statistical analyses are reported in the Table 2b.

**TABLE 1.** GLMM output testing the effect of the interaction between female genotype and (normalized) weight on the number of males attracted in the enclosure setup. Significant p-values are highlighted in bold.

| Group | Name | Variance | SD |  |
| --- | --- | --- | --- | --- |
| Replicate | (intercept) | 0.16 | 0.40 |  |
| Trap position | (intercept) | 0.89 | 0.94 |  |
| Female ID | (intercept) | 0.37 | 0.61 |  |
| Fixed effects: |  |  |  |  |
|  | Estimate | SE | z | P |
| Intercept (Wy) | -3.87 | 0.49 | -7.95 | **1.87e-15** |
| WW | -0.38 | 0.45 | -0.84 | 0.40 |
| yy | 0.63 | 0.41 | 1.55 | 0.12 |
| Weight | -0.79 | 0.35 | -2.24 | **0.03** |
| WW genotype * weight | 0.99 | 0.45 | 2.21 | **0.03** |
| yy genotype * weight | 1.88 | 0.59 | 3.19 | **0.001** |

**TABLE 2.**
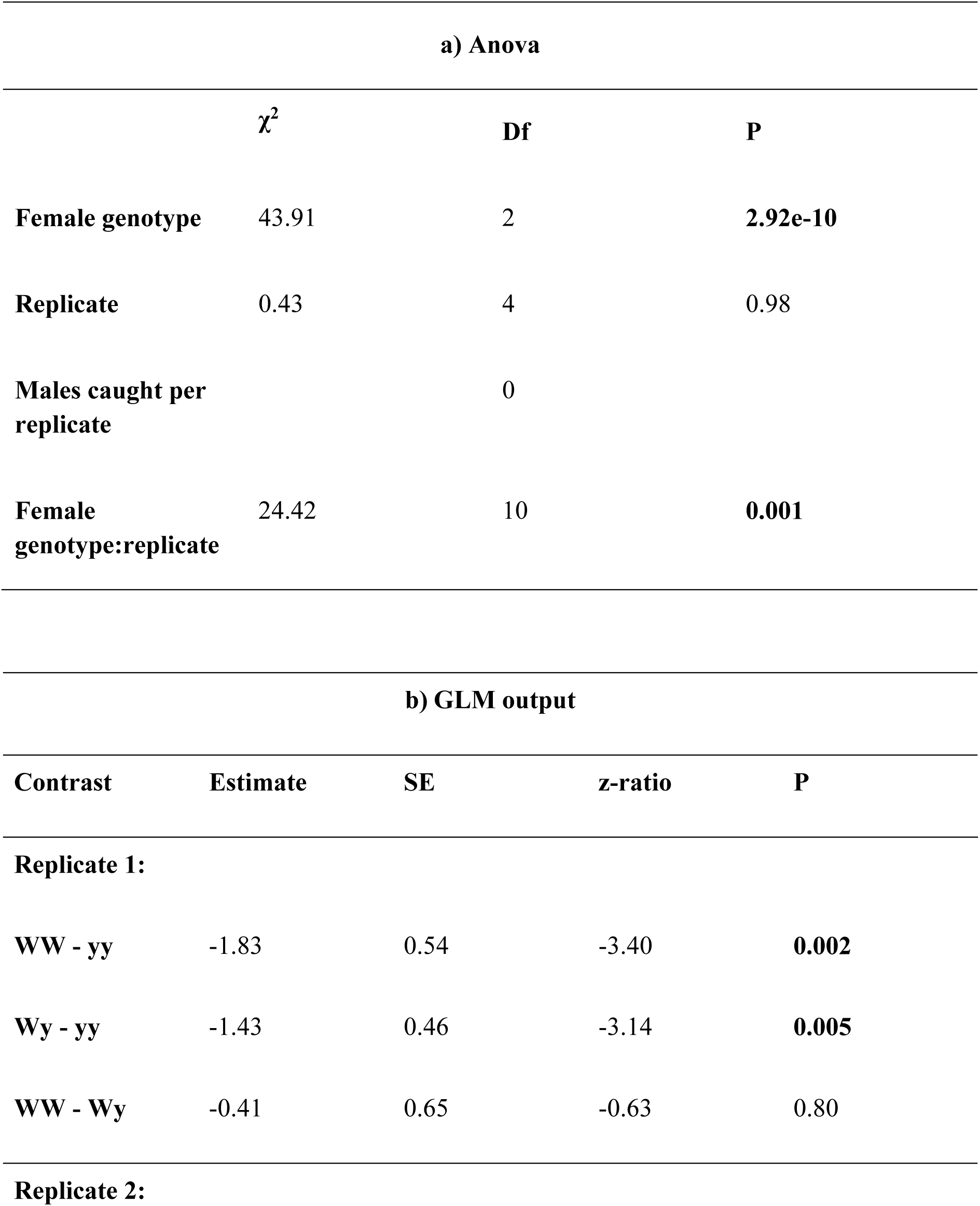
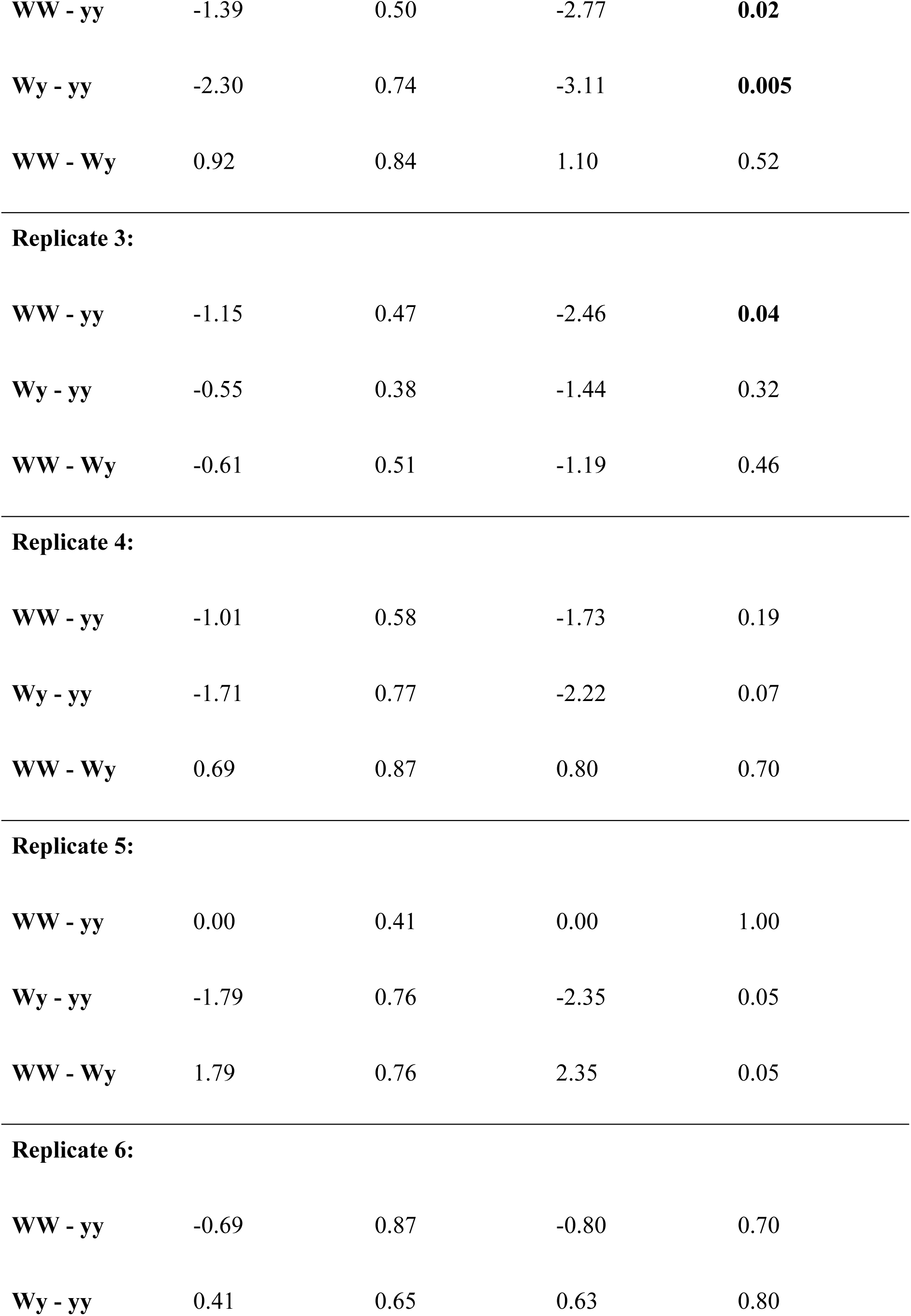
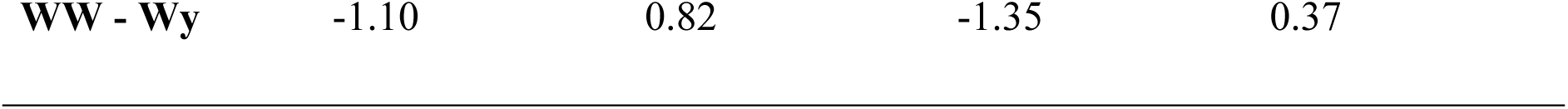
a) Overall effect of the fixed factors tested with Chi-square for the daily number of males attracted by female genotype in the enclosure setup. b) Pairwise comparison based on estimated marginal means (EMM) based on the GLM output on the number of males attracted by night in the enclosure experiment. Replicates refer to the six nights of experiments. Significant differences in the number attracted are marked in bold.

| a) Anova |  |  |  |
| --- | --- | --- | --- |
| | $\chi^2$ | Df | P |
| Female genotype | 43.91 | 2 | **2.92e-10** |
| Replicate | 0.43 | 4 | 0.98 |
| Males caught per replicate |  | 0 |  |
| Female genotype:replicate | 24.42 | 10 | **0.001** |

| b) GLM output |  |  |  |  |
| --- | --- | --- | --- | --- |
| Contrast | Estimate | SE | z-ratio | P |
| Replicate 1: |  |  |  |  |
| WW - yy | -1.83 | 0.54 | -3.40 | **0.002** |
| Wy - yy | -1.43 | 0.46 | -3.14 | **0.005** |
| WW - Wy | -0.41 | 0.65 | -0.63 | 0.80 |

Replicate 2:
|  |  |  |  |  |
| --- | --- | --- | --- | --- |
| **WW - yy** | -1.39 | 0.50 | -2.77 | **0.02** |
| **Wy - yy** | -2.30 | 0.74 | -3.11 | **0.005** |
| **WW - Wy** | 0.92 | 0.84 | 1.10 | 0.52 |

**Replicate 3:**
|  |  |  |  |  |
| --- | --- | --- | --- | --- |
| **WW - yy** | -1.15 | 0.47 | -2.46 | **0.04** |
| **Wy - yy** | -0.55 | 0.38 | -1.44 | 0.32 |
| **WW - Wy** | -0.61 | 0.51 | -1.19 | 0.46 |

**Replicate 4:**
|  |  |  |  |  |
| --- | --- | --- | --- | --- |
| **WW - yy** | -1.01 | 0.58 | -1.73 | 0.19 |
| **Wy - yy** | -1.71 | 0.77 | -2.22 | 0.07 |
| **WW - Wy** | 0.69 | 0.87 | 0.80 | 0.70 |

**Replicate 5:**
|  |  |  |  |  |
| --- | --- | --- | --- | --- |
| **WW - yy** | 0.00 | 0.41 | 0.00 | 1.00 |
| **Wy - yy** | -1.79 | 0.76 | -2.35 | 0.05 |
| **WW - Wy** | 1.79 | 0.76 | 2.35 | 0.05 |

**Replicate 6:**
|  |  |  |  |  |
| --- | --- | --- | --- | --- |
| **WW - yy** | -0.69 | 0.87 | -0.80 | 0.70 |
| **Wy - yy** | 0.41 | 0.65 | 0.63 | 0.80 |
| **WW - Wy** | -1.10 | 0.82 | -1.35 | 0.37 |

##### Strategies of female attractiveness

Within night (of the enclosure setup), yy females attracted males significantly earlier than WW (coxph; coef = −1.13 ± 0.43, z = −4.38, *P* < 0.001) and Wy females (coxph; coef = −0.91 ± 0.42, z = −2.21, *P* = 0.03) (Fig. 4). In all nights of the enclosure setup, males were first found in traps baited with a yy female. In only two instances, males were also found in a trap baited with a WW (1/6 replicate) and a Wy female (1/6 replicate) (Fig. 4). When we conducted tests to examine whether the time at which males were initially found in the trap or the duration of female activity could explain the number of males attracted within a single night, our findings revealed that a higher proportion of males were attracted by females that initiated their attraction behavior earlier in the night (χ^2^ = 6.95, df =1, *P* = 0.008) but was not affected by the length of time a female attracted males for throughout the night (χ^2^ = 2.59, df = 1, *P* = 0.11; GLMM: estimate = 0.39 ± 0.25, z = 1.61, P = 0.11) or by the number of males left in the tent (χ^2^ = 2.34, df = 1, P = 0.13). The later the female would start attracting, the lower the proportion of males caught (GLMM; estimate = −0.63 ± 0.24, z = −2.64, *P* = 0.008) (Random effects; trap position: variance = 0.20, SD = 0.45, replicate: variance = 0.09, SD = 0.30) (Fig. 4).

**FIGURE 4.**
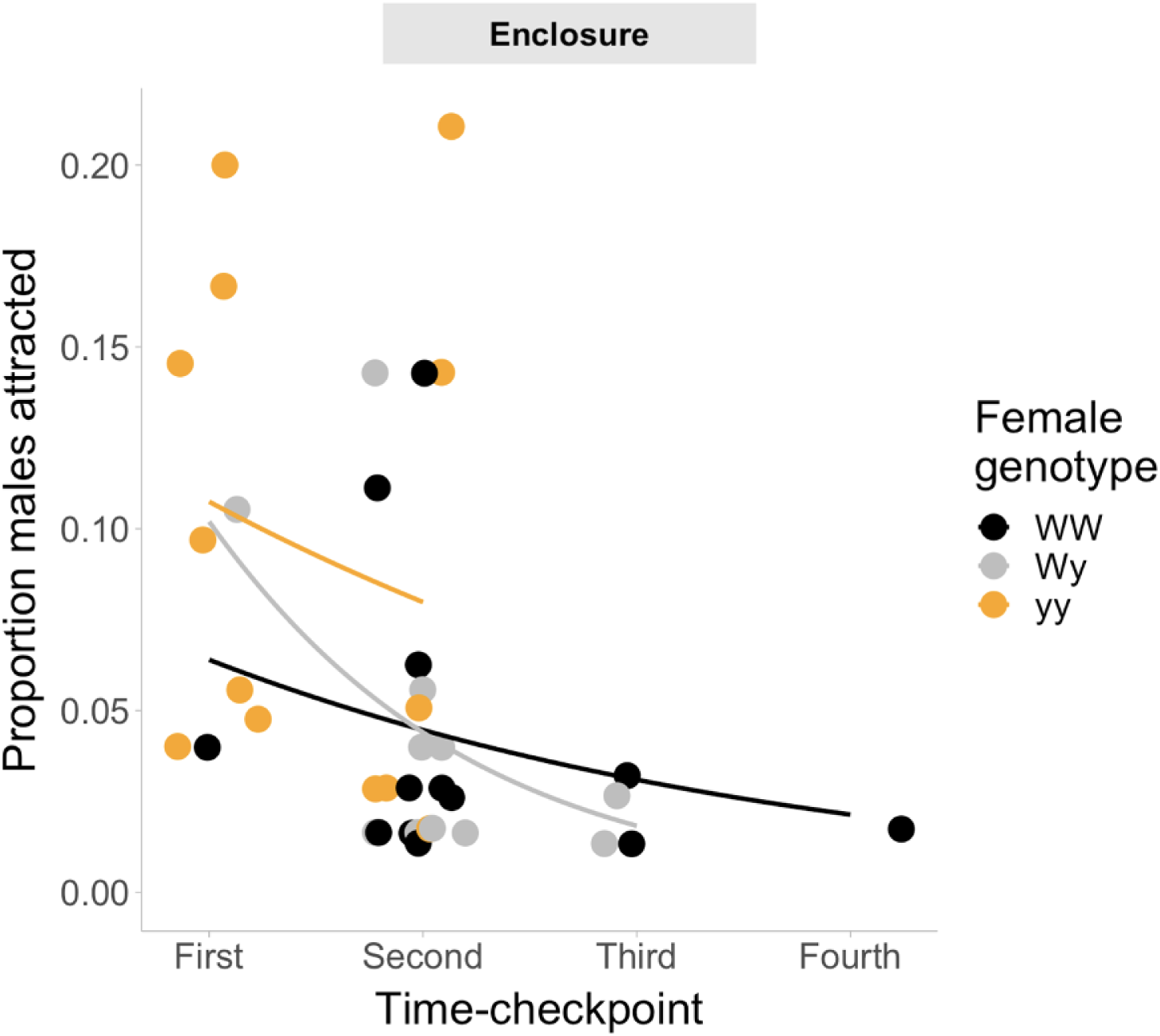
Proportion of males attracted by female according to the time-checkpoint the first male was found in the trap. ‘First’ refers to the first time-checkpoint a male was found in traps, and so on until the fourth within night. This figure also shows that, typically, yy females are more likely to attract males earlier than WW and Wy females.

### Male order of arrival

#### Antennal morphology

Overall, we observed no connection between color variation and antennal structure among wild males. When it comes to males raised in the laboratory, the colour genotype had no discernible impact on antennal length and lamellae count. However, WW males exhibited broader antennae compared to Wy and yy males. Importantly, we found that there were no distinctive differences in antennal structure based on morph between lab-reared and wild males, excluding the possibility of laboratory selection on male antennal morphology. For a more comprehensive breakdown of the results, please refer to the Supporting Information section “Further details of the results on genotype-based and lab-reared vs wild caught differences in male antennal morphology”.

#### Effect of antennal morphology and genetic color morphs on the order of arrival

In the field setup, antennal length (LR χ^2^ = 0.08, df = 1, P = 0.78), area (LR χ^2^ = 0.73, df = 1, P = 0.39) and number of lamellae (LR χ^2^ = 0.04, df = 1, P = 0.85) did not affect the order of the male arrival to the females. In the enclosure, while antennal length (LR χ^2^ = 0.86, df = 1, P = 0.35) and area (LR χ^2^ = 2.31, df = 1, P = 0.13) did not affect the order of male arrival to the traps, we found a significant effect of the number of lamellae (LR χ^2^ = 6.13, df = 1, P = 0.01). Males with a higher number of lamellae reached the females faster than males with a lower number of lamellae (coeff = 0.07 ± 1.07, z = 2.47, P = 0.01) (Fig. S3). Notably, males captured later in the enclosure had, on average, fewer lamellae than males attracted over the first two nights (72 ± 1.31 SE vs 77 ± 0.58 SE) (Fig. S3). Male weight (LR χ^2^ = 0.87, df = 1, P = 0.35) and age (LR χ^2^ = 0.001, df = 1, P = 0.96) did not affect arrival to females in the enclosure setup. It was interesting to notice that male morph (in the field setup; LR χ^2^ = 5.34, df = 1, P = 0.02) and male genotype (in the enclosure setup; LR χ^2^ = 8.40, df = 2, P = 0.02) significantly affected the male order of arrival to females. In the field, yellow males were more likely to arrive earlier than white males (coeff = 0.42 ± 0.18, z = 2.35, P = 0.02) (Fig. 5a). In the enclosure, yy males instead took longer to reach females compared to the other two genotypes (yy vs WW; coeff = 0.76 ± 0.29, z = 2.65, P = 0.008; yy vs Wy; coeff = 0.62 ± 0.27, z = 2.28, P = 0.02) (Fig. 5b, Table S7b). Finally, we found that the number of antennal lamellae did not differ between males attracted by females of the three different genotypes of the enclosure setup (F_(2,117)_= 1.53, P = 0.22).

**FIGURE 5.**
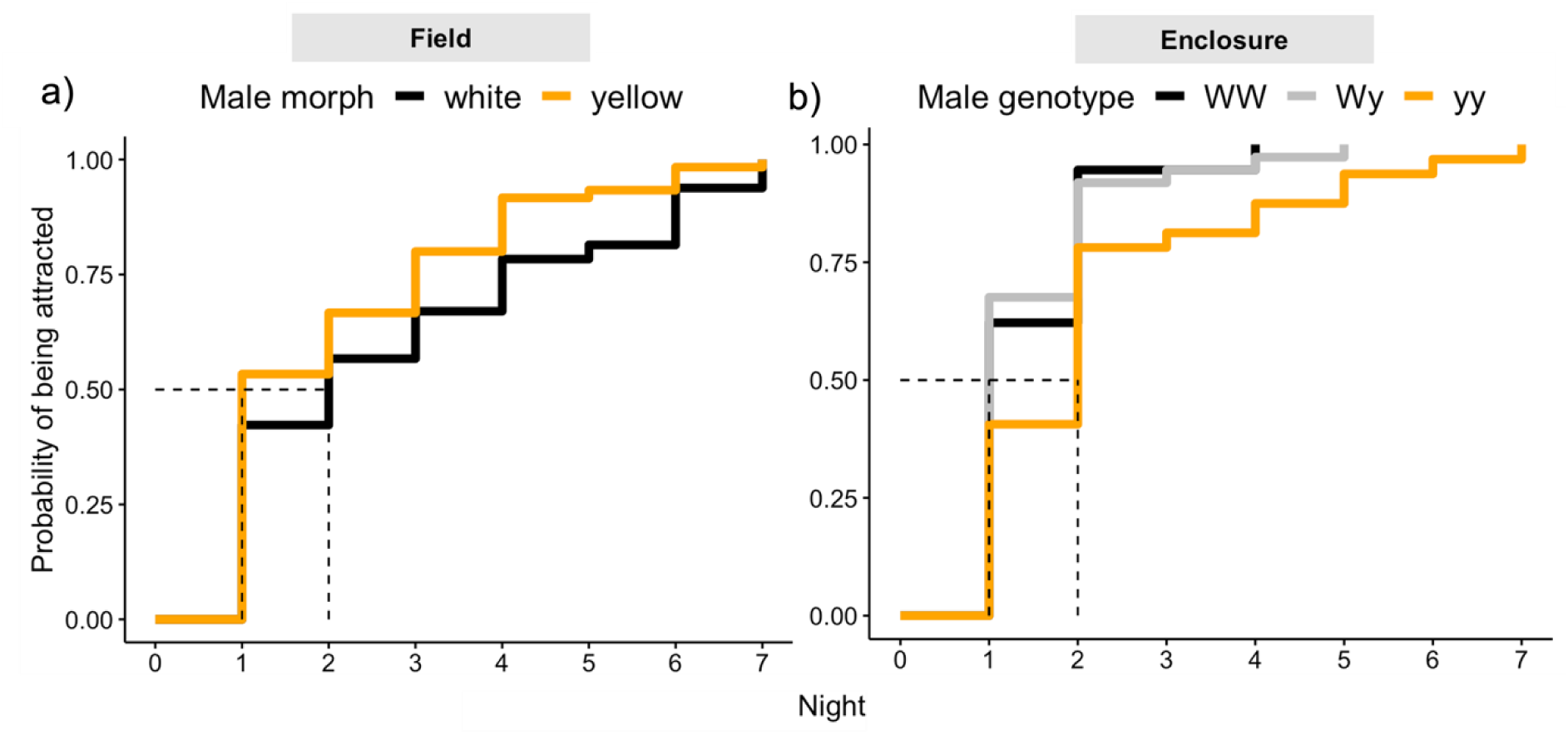
The order of white and yellow male’s arrival was affected by the experimental setup. a) In the field, yellow males reached the females faster than their counterpart. b) In the enclosure, white (WW and Wy) males were faster in reaching the females compared to yellow (yy) males.

## Discussion

The persistence of phenotypic and genetic variation in wild populations is often the result of complex interactions between multiple mechanisms and selective pressures. This is especially true for species characterized by discrete color morphs, as these morphs often differ in multiple sets of traits (Fisher, 1930; Sinervo & Lively, 1996). Such variation can, in part, be attributed to pleiotropic effects of the color locus on behavior, physiology, and life-history traits, influencing intrapopulation dynamics and individual interactions. Within the context of precopulatory selection, this variation becomes particularly relevant when considering how morph-linked traits and variation in chemical communication channels affect mate choice. In this study, we aim to understand how variation in sexual traits, both in signalers and receivers, impact mate responses, mate choices, and ultimately contribute to the maintenance of genetic diversity in reproductive traits. This research sheds light on a potential mechanism for the persistence of polymorphisms under precopulatory selection. In summary, we found that heavier Wy females attracted a lower proportion of males compared to WW and yy females, at least in the high density setup. yy females were generally more attractive than the other two genotypes, likely due to earlier pheromone release during the night. Males with a higher number of lamellae and white males reached females faster in the high density setup, whereas yellow males located females faster than white males in the low density setup.

We found, unexpectedly, that yy females were more attractive to males compared to WW and Wy females, likely due to females’ earlier calling strategy as traps baited with yy females were more likely to collect males earlier than the traps baited with WW and Wy females. In turn, male color genotype and antennal morphology affected males’ ability to reach females. Yellow males reached females faster in the field, while white males, and males with higher number of antennal lamellae located females faster in the enclosure. These results show that variation in sexual signals, male antennae, and morph-specific precopulatory strategies can affect mate choice and recruitment.

Contrary to our prediction of correlation between female attractiveness and reproductive success, we found that heavier Wy females attracted a lower proportion of males compared to WW and yy females, at least in the enclosure setup. Specifically, an increase in weight corresponded to a decreased Wy but increased yy female attractiveness. yy females were in general more attractive than others early in the night which was likely due to females’ earlier calling strategy as traps baited with yy females were more likely to collect males earlier than the traps baited with WW and Wy females. It should be noted, however, that in the field setup, the power of the test was lower due to the smaller sample size. Associations between color locus and female attractiveness have also been extensively reported in *Colias* butterflies, with ‘alba’ (A-genotype) females typically being more attractive than ‘non-alba (aa genotype) females (Graham et al., 1980; Kemp & Macedonia, 2007). The opposing effect of weight for Wy and yy females may reflect genotype-specific resource allocation strategies during larval development that would be typical of a capital breeding species, such as the wood tiger moth. While more research is needed to explore this further, it is possible that Wy females invest more resources during development to secure reproductive output while yy females may allocate more resources into pheromone production. This suggests that WW and Wy females may face a higher risk of remaining unmated compared to yy females, at least when population density is low. De Pasqual et al., (2022) recently found that Wy females have higher fertility, offspring survival and egg hatching success compared to either homozygote whereas yy females had a significantly higher probability of failing to have offspring compared to W-females, as they had an increased likelihood in failing to lay viable eggs. Increased attractiveness may be advantageous for yy females as it provides more opportunities to encounter several males, potentially enhancing their reproductive success by increasing the likelihood of pairing with an optimal male among multiple competitors (reviewed in Wong & Candolin, 2005). For example, in the three-spined stickleback (*Gasterosteus aculeatus*), female choice for males with higher parental abilities is facilitated by male competition because it increases sexual signaling expression in dominant males (Candolin, 1999). Males attracted to yy females may, instead, incur the risk of reproductive failure, thereby generating potential sexual conflict in this study system. We should, however, note that yy females’ higher attractiveness is typically limited to few individuals, suggesting that attracting many males may still be costly for females as they may experience strong male harassment.

Signalers’ attractiveness can also change according to population density and the progression of the season (Evenden et al., 2015; Niemelä et al., 2021). The higher yy female attractiveness across multiple nights in the enclosure, suggests that yy females may be more successful when males are present in higher densities. Population density can indeed affect individual attractiveness and vary over time, as recently shown for male crickets (*Gryllus campestris*) (Niemelä et al., 2021). In addition, environmental factors can also affect female attractiveness (Groot, 2014). In the enclosure setup, yy female advantage declined when the night temperature was unusually high. The daily temperature during the third male replicate was about 9°C higher than the previous nights (∼29°C vs ∼20.5°C). Temperature can indeed affect sexual activity in moths (Webster & Cardé, 1982; Groot, 2014) with an increase in temperature causing substantial delay in the onset of calling (Schal & Cardé, 1986), decrease the pheromone titer (Giebultowicz et al., 1992), or make male and female sexual activities asynchronous (Giebultowicz et al., 1992).

Moreover, variation in female attractiveness can also result from different signaling strategies (Umbers et al., 2015), including a tendency of early-life (Baudry et al., 2021) or later-in-life (Pham et al., 2021; Umbers et al., 2015) signaling investment, manifested with increased signaling/calling effort either soon after eclosion (Baudry et al., 2021) or as females age (Pham et al., 2021; Umbers et al., 2015). In all six nights of the enclosure experiment, males were found first in traps baited with a yy female and the proportion of males attracted by females was higher the earlier males were found in traps. Therefore, we propose that yy female attractiveness may benefit from a strategy of early night calling and a wider overlap with male searching activity, although this remains to be tested. The early calling strategy of yy females may have a particular advantage in high density populations, which would explain their constant advantage in the enclosure environment.

Moth mating systems are characterized by scramble competition, in which male mating success depends on their ability to locate females efficiently. Scrambling males compete by using traits that allow them to quickly reach females (e.g., larger wingspan) or through effective mate-location structures (e.g., antennae) (Herberstein et al., 2017). Contrary to our expectation that, irrespective of the experimental setup, white males would reach females faster than yellow males, we found that the order of male arrival was context-specific (i.e., different between field and enclosure setup). White males reached females faster than the other male morph in the enclosure, whereas yellow males reached females faster than whites in the field. Therefore, white males may succeed better when the population density is high (i.e., the enclosure setup) whereas yellow males may succeed better in low population density (i.e., the field setup). Previous research also showed context-dependent morph-differences; white and yellow males are characterized by morph-specific flight activities, with yellow males having a narrower daily activity pattern than white males (Rojas et al., 2015) while a mark-recapture study found that yellow males disperse less than whites (Gordon et al., unpublished). The opposite morph-arrival patterns of the two experiments therefore suggests that male behavior may be affected by morph-specific behavioral strategies, antennal morphology (see next paragraph), and population density. While this may hint to either a role of intrasexual competition due to different male densities or to male’s behavioral strategies, further investigation is needed to determine which of the described factors can explain our results.

In addition to color morph-specific differences in the order of male arrival to traps, males with denser (i.e., higher lamellae count) antennae reached females faster in the enclosure setup than males with a lower number of lamellae. Since we can exclude the possibility that this result was driven by laboratory adaptation, the number of lamellae is likely a trait that benefits males in locating females, particularly in higher density populations. This trait may be under sexual selection as it improves pheromone perception and may be useful to outcompete other males. In this species, male-male competition is notably intense due to the male-biased operational sex-ratio prevalent in the field. In field observations multiple males competing for and courting the same female can frequently be observed. Additionally, most females engage in mating only once. Furthermore, in cases where females do engage in multiple matings, there exist no distinct advantage for either the last or first male involved (Santostefano et al., 2018). In species in which mate recruitment is initiated through the release of long-range sex pheromone, such as in moths, females are likely to play an important role in the first stage of attraction. In both experimental setups, females likely released pheromone for longer during the night than they naturally would, as individuals were not allowed to engage in copulation. This allowed us to test for potential disassortative attractiveness, a mechanism that favors the maintenance of both color alleles in the population under balancing selection, as shown for color coat in wolves (Hedrick et al., 2016) and white-throated sparrows (Tuttle et al., 2016). Although not in the strict sense of the genotype (e.g., yy females to attract more WW or Wy males), we may have found evidence for some form of disassortative attractiveness in the enclosure experiment. Since white (WW and Wy) males reached females faster than yy males, and typically, yy females attracted males earlier than the other two genotypes, highly competitive environments may favor the coexistence of both color alleles.

To conclude, we did not find evidence of assortative or disassortative attraction but we found that morph-linked behaviour and reproductive traits affect individual precopulatory success, allowing us to expand our understanding on how complex polymorphisms can affect between-individual interactions in nature. These precopulatory strategies linked to color morphs may, at least in part, be determined by pleiotropic effects of the color locus on sexual traits (e.g., female sex pheromones and male receiving structures). A parallel can be drawn with *Drosophila melanogaster*, in which the lack of the *yellow* gene function, disrupts the expression of body pigmentation, male courtship behavior and mating success (Bastock, 1956; Massey et al., 2019; Wilson et al., 1976). Our study represents a rare case of interplay between morph-related precopulatory strategies and variation in chemical communication channels which, altogether, may offer an additional mechanism for the maintenance of intrapopulation phenotypic variation under precopulatory selection.

## Acknowledgments

We thank Sara Calhim and Janne Valkonen for help with statistical analyses, and Cristina Ottocento, Sandra Winters, Johan Fitzpatrick, Maya Evenden, and James Barnett for helpful comments and discussion on a previous version of the manuscript. This study was funded by the Academy of Finland (project no. 345091) to JM and by the Finnish Cultural Foundation (grant #00220196) to CDP.

## Data availability statement

Data will be made available upon manuscript acceptance on the Dryad Digital Repository

