## Supplementary material for "Morph-linked variation in female pheromone signaling and male response in a polymorphic moth": Fig. S1, Fig. S2, Table S1, Table S2, Table S3, Table S4, Table S5, Table S6, Fig. S3

**SUPPORTING INFORMATION**

**Further details on the mating scheme and rearing protocol (referred to Materials and methods section on the main text)**

The laboratory stock population was established in 2013 from wild-caught individuals collected across the different Finnish sub-populations (visit laji.fi for species distribution mapping). New individuals are introduced yearly to the stock to preserve the genetic variability of the laboratory population. To ensure offspring paternity, during stock maintenance, one female and one male of known color genotype are paired in transparent mating boxes (13 H x 7 W x 9 L cm) provided with mesh on the lid for aeration. The mating scheme included matings between all genotype combinations by selecting genotyped individuals from as many families as available in the stock. Once mated, females lay, on average, 250 eggs which hatch into larvae after approximately seven days (Chargé et al., 2016). Larvae are polyphagous and feed on a variety of weedy plants (e.g., *Plantago* sp., and *Taraxacum* sp.,). Fourteen days after hatching, larvae are divided in rearing containers of 30 larvae each, where they grew until pupation. The laboratory stock is maintained in a greenhouse that roughly followed the outdoor temperature (20 – 25 °C) and the natural light.

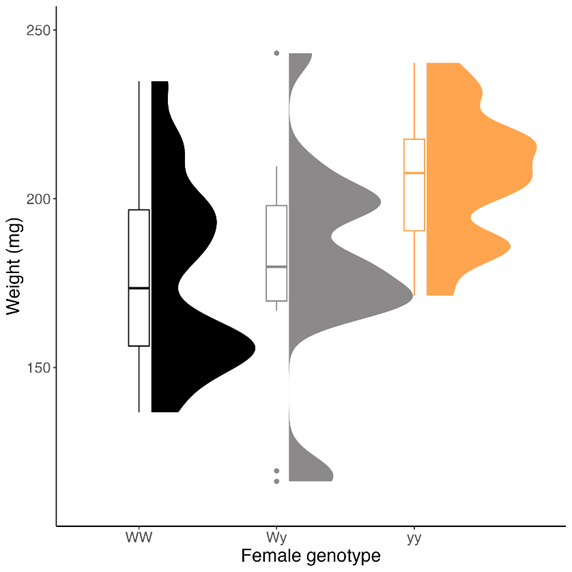

FIGURE S1	Graph reporting the distribution of the female weight (y-axis) for each genotype (x-axis) for the females tested in the enclosure setup. The upper and lower borders of the boxplots indicate the first and third quartile. The thick bar within the box indicates the group median. Whiskers above and below each box extend to a maximum of 1.5 times the interquartile range. Outliers are indicated with circles. The raincloud plot (on the right of each boxplot) shows the half-density distribution of values by group (i.e., genotype).

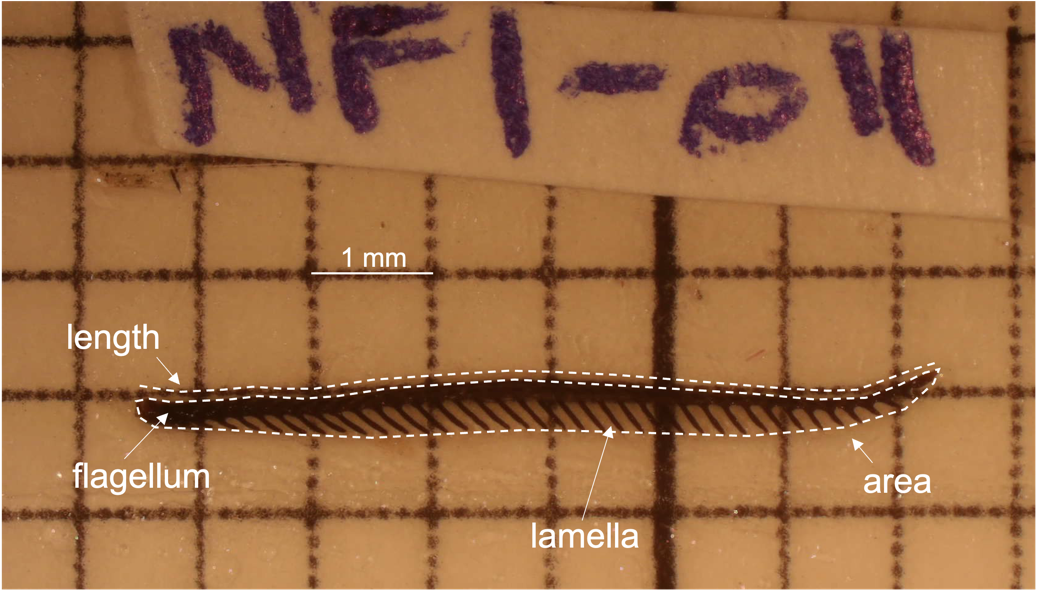

FIGURE S2	Male’s ability to locate females was linked to antennal morphology. Once the antenna was excised from the male’s head, antennal measurements were obtained through photography (1 x magnification). The area was measured by including the flagellum and one side of the lamellae. The length was measured as the length of the flagellum. The number of lamellae were counted manually.

Further details on the model selection based on AIC (referred to the Statistical analyses section)

TABLE S1	Each table reports model selection based on the AIC throughout the analyses section. The selected model is highlighted in bold.

a) Female attractiveness - Field setup

|  | model | df | AIC |
| --- | --- | --- | --- |
| Likelihood of attractiveness | genotype * night + weight + (1\|night:location) | 8 | 90.99 |
|  | **genotype + night + weight + (1\|night:location)** | **6** | **90.12** |
| Number of males attracted | **genotype + night + weight + (1\|night:location)** | **7** | **244.83** |

*b) Female attractiveness - Enclosure setup*

|  | model | df | AIC |
| --- | --- | --- | --- |
| Likelihood of attractiveness | genotype * weight + (1\|trap position) + (1\|replicate) | 9 | 74.88 |
|  | **genotype + weight + (1\|trap position) + (1\|replicate)** | **7** | **75.40*** |
| Number of males attracted | **genotype * weight + (1\|trap position) + (1\|replicate)** | **9** | **222.49** |
|  | genotype + weight + (1\|trap position) + (1\|replicate) | 7 | 227.55 |
| Disassortative attractiveness | Female genotype * male genotype + males left in the tent + (1\|femaleID) | - | - |

*** Given that the difference in AIC is lower than 2, we retained the simpler model (i.e., without the interaction) following “Burnham, K. P., Anderson, D. R. (1998). Practical use of the information-theoretic approach. In Model selection and inference: A practical information-theoretic approach, pp. 75-117, Springer”.**

*c) Temporal variation in female attractiveness*

|  | **Df** | **AIC** |
| --- | --- | --- |
| onset * duration of pheromone release + males left in the tent | 7 | 185.46 |
| **onset + duration of pheromone release + males left in the tent** | **6** | **185.58*** |

*d) Effect of antennal morphology and genetic color morph on the order of arrival - field setup*

|  | **Df** | **AIC** |
| --- | --- | --- |
| length * male morph + area * male morph + lamellae * male morph | 7 | 1105.84 |
| length * male morph + area + lamellae + male morph | 5 | 1101.93 |
| length + area * male morph + lamellae + male morph | 5 | 1102.17 |
| length + area + lamellae * male morph + male morph | 5 | 1102.44 |
| **length + area + lamellae + male morph** | **4** | **1100.46** |

*e. Effect of antennal morphology and genetic color morph on the order of arrival - enclosure setup*

|  | **Df** | **AIC** |
| --- | --- | --- |
| length * male genotype + area * male genotype + lamellae * male genotype + male weight + male age | 13 | 736.11 |
| length * male genotype + area + lamellae + male weight + male age | 9 | 736.98 |
| length + area * male genotype + lamellae + male weight + male age | 9 | 736.47 |
| length + area + lamellae * male genotype + male weight + male age | 9 | 734.06 |
| **length + area + lamellae + male genotype + male weight + male age** | **7** | **733.48** |

Table S2 - Model output for the models testing female attractiveness in the field setup

a) Likelihood of female attractiveness

| Group | Name | Variance | SD |  |
| --- | --- | --- | --- | --- |
| Night_location | (intercept) | 0.73 | 0.85 |  |
| Fixed effects: |  |  |  |  |
|  | Estimate | SE | z | P |
| Intercept (yy) | 0.98 | 0.88 | 1.12 | 0.263 |
| WW | 0.22 | 0.74 | 0.30 | 0.77 |
| Wy | 0.28 | 0.75 | 0.37 | 0.71 |
| Weight | 0.33 | 0.33 | 1.01 | 0.31 |
| Night | -0.14 | 0.18 | -0.77 | 0.44 |

b) Number of males attracted

| **Group** | **Name** | **Variance** | **SD** |  |
| --- | --- | --- | --- | --- |
| **Night:location** | (intercept) | 0.52 | 0.72 |  |
| **Fixed effects:** |  |  |  |  |
|  | **Estimate** | **SE** | **z** | **P** |
| **Intercept (yy and night 1)** | **1.91** | **0.41** | **4.61** | **4.08e-06** |
| **WW** | -0.15 | 0.26 | -0.60 | 0.55 |
| **Wy** | -0.38 | 0.22 | -1.74 | 0.08 |
| **Night 2** | **-1.27** | **0.58** | **-2.16** | **0.03** |
| **Night 3** | **-1.50** | **0.59** | **-2.55** | **0.01** |
| **Night 4** | **-1.60** | **0.58** | **-2.77** | **0.01** |
| **Night 5** | **-2.37** | **0.79** | **-2.99** | **0.003** |
| **Night 6** | **-1.20** | **0.62** | **-1.95** | **0.05** |
| **Night 7** | **-2.12** | **0.68** | **-3.10** | **0.002** |
| **Female weight** | 0.32 | 0.17 | 1.87 | 0.06 |

**c) Daily number of males attracted by female genotype**

|  | **Estimate** | **SE** | **z** | **P** |
| --- | --- | --- | --- | --- |
| **Intercept (WW and night 2)** | **0.81** | **0.33** | **2.43** | **0.02** |
| **Wy** | 0.39 | 0.46 | 0.86 | 0.39 |
| **yy** | -0.81 | 0.67 | -1.22 | 0.22 |
| **Night 3** | 0.04 | 0.50 | 0.07 | 0.94 |
| **Night 4** | 0.20 | 0.45 | 0.45 | 0.66 |
| **Males caught by night** | NA | NA | NA | NA |
| **Wy:night3** | -0.73 | 0.74 | -0.98 | 0.33 |
| **yy:night3** | 0.37 | 0.87 | 0.43 | 0.67 |
| **Wy:night4** | -1.00 | 0.68 | -1.46 | 0.14 |
| **yy:night4** | -0.49 | 0.93 | -0.52 | 0.60 |

**d) Disassortative attractiveness**

| **Group** | **Name** | **Variance** | **SD** |  |
| --- | --- | --- | --- | --- |
| **Female ID** | (intercept) | 0.64 | 0.80 |  |
| **Night_location** | (intercept) | 0.65 | 0.81 |  |
| **Fixed effects:** |  |  |  |  |
|  | **Estimate** | **SE** | **z** | **P** |
| **Intercept (WW and white morph)** | -0.47 | 0.3 6 | -1.31 | 0.19 |
| **Wy** | -0.07 | 0.46 | -0.16 | 0.87 |
| **yy** | 0.41 | 0.43 | 0.94 | 0.35 |
| **Yellow morph** | -0.53 | 0.35 | -1.50 | 0.13 |
| **Wy:yellow morph** | 0.53 | 0.47 | 1.13 | 0.26 |
| **yy:yellow morph** | -0.24 | 0.45 | -0.54 | 0.59 |

TABLE S3	Summary table of female genotypes’ attractiveness as an overview of the disassortative attractiveness. The left side of the table reports the number of males per morph, and their sum, caught in the field. The right side of the table reports the number of males per genotype caught in the enclosure.

|  | **Males attracted by females tested in the field setup** | | | **Males attracted by females tested in the enclosure** | | | |
| --- | --- | --- | --- | --- | --- | --- | --- |
| **Female genotype** | **White males** | **Yellow males** | **All males** | **WW males** | **Wy males** | **yy males** | **All**  **males** |
| WW | 22 | 13 | 35 | 9 | 11 | 8 | 28 |
| Wy | 21 | 21 | 42 | 7 | 6 | 11 | 24 |
| yy | 41 | 19 | 60 | 28 | 28 | 21 | 77 |
| **Total** | **84** | **53** | **137** | **44** | **45** | **40** | **129** |

Table S4 - Model output for the models testing female attractiveness in the enclosure setup

a) **Likelihood of female attractiveness**

| **Group** | **Name** | **Variance** | **SD** |  |
| --- | --- | --- | --- | --- |
| **Female ID** | (intercept) | 0 | 0 |  |
| **Trap position** | (intercept) | 0.99 | 0.99 |  |
| **Replicate** | (intercept) | 0 | 0 |  |
| **Fixed effects:** |  |  |  |  |
|  | **Estimate** | **SE** | **z** | **P** |
| **Intercept (yy)** | 1.19 | 0.74 | 1.60 | 0.11 |
| **Wy** | -0.41 | 0.91 | -0.45 | 0.65 |
| **WW** | -0.18 | 0.91 | -0.20 | 0.84 |
| **Weight** | -0.04 | 0.39 | -0.11 | 0.91 |

b) **Disassortative attractiveness**

| **Group** | **Name** | **Variance** | **SD** |  |
| --- | --- | --- | --- | --- |
| **Female ID** | (intercept) | 1.13 | 1.06 |  |
| **Replicate** | (intercept) | 0 | 0 |  |
| **Fixed effects:** |  |  |  |  |
|  | **Estimate** | **SE** | **z** | **P** |
| **Intercept (WW male and WW male)** | **-3.91** | **0.48** | **-8.19** | **2.62e-16** |
| **Wy female** | -0.24 | 0.69 | -0.34 | 0.73 |
| **yy female** | 1.00 | 0.59 | 1.68 | 0.09 |
| **Wy male** | 0.25 | 0.47 | 0.53 | 0.60 |
| **yy male** | -0.16 | 0.50 | -0.32 | 0.75 |
| **Wy female:Wy male** | -0.38 | 0.74 | -0.52 | 0.61 |
| **yy female:Wy male** | -0.21 | 0.55 | -0.38 | 0.71 |
| **Wy female:yy male** | 0.43 | 0.72 | 0.60 | 0.55 |
| **yy female:yy male** | -0.19 | 0.59 | -0.33 | 0.74 |

**Further details on the statistical analyses performed on male antennal morphology (referred to the Material and methods section in the main text)**

Before delving into the potential role of male trait variation in affecting the order of arrival to traps, we checked for potential differences in antennae due to color genotype, weight, and male origin (i.e., wild-caught or lab-reared). We thus tested whether antennal length, area, and number of lamellae (response variables) were affected by genotype or weight (lab-reared males) or color morph (wild-caught males) (fixed factors). We set two linear models for antennal length and area, while the number of lamellae was modelled with Poisson distribution. Furthermore, because the antenna analyses were performed on wild- and lab-reared males, we compared antennal morphology to disentangle any difference due to laboratory rearing. We thus compared the antennal length, area, and number of lamellae (response variables) by setting an interaction between male origin (i.e., wild-caught or lab-reared) and male morph (fixed factor). This allowed us to also test for potential within-morph differences between wild- and lab-reared males. We set four linear models, two for antennal length and two for antennal area, comparing the model with and without interaction. For the number of lamellae, we set a General Linear Model (GLM) with the response variable modelled with Poisson distribution We chose the model based on the lowest AIC (Table S5). In addition, we tested for collinearity between the three antennal traits through the Variance Inflation Factor (VIF) (Table S6).

TABLE S5	The table shows the model selection based on the lowest AIC when testing for differences in antennal morphology due to the origin of the males

| Antennal area | Df | AIC |
| --- | --- | --- |
| Male origin * male morph | 5 | 41.07 |
| Male origin + male morph | 4 | **40.84** |
| Antennal length | Df | AIC |
| Male origin * male morph | 5 | 223.26 |
| Male origin + male morph | 4 | **221.45** |
| Lamella number | Df | AIC |
| Male origin * male morph | 4 | 1699.28 |
| Male origin + male morph | 3 | **1697.62** |

TABLE S6	The table shows the VIF coefficient between the three antennal traits, i.e., length, area and number of lamellae.

Field setup:

| Trait -> | Length | Area | Lamellae | Male morph |
| --- | --- | --- | --- | --- |
| VIF | 2.1 | 2 | 1.1 | 1.01 |

**Enclosure setup:**

| **Trait ->** | **Length** | **Area** | **Lamellae** | **Male genotype** | **Male weight** | **Male age** |
| --- | --- | --- | --- | --- | --- | --- |
| **VIF** | 1.7 | 1.8 | 1 | 1.1 | 1.2 | 1 |

Table S7 - Models output for the models testing the effect of antennal morphology and genetic color morphs on the order of male arrival

a) Field setup

|  | **Coef** | **SE(coef)** | **z** | **P** |
| --- | --- | --- | --- | --- |
| **Length** | -0.10 | 0.37 | -0.28 | 0.78 |
| **Area** | -0.42 | 0.49 | -0.85 | 0.39 |
| **Lamellae** | 0.00 | 0.02 | 0.18 | 0.85 |
| **yellow males** | **0.42** | **0.18** | **2.35** | **0.02** |

b) Enclosure setup

|  | **Coef** | **SE(coef)** | **z** | **P** |
| --- | --- | --- | --- | --- |
| **Length** | 0.45 | 0.49 | 0.93 | 0.35 |
| **Area** | -1.02 | 0.68 | -1.50 | 0.13 |
| **Lamellae** | **0.07** | **0.03** | **2.47** | **0.01** |
| **WW males** | **0.76** | **0.29** | **2.65** | **0.01** |
| **Wy males** | **0.62** | **0.27** | **2.28** | **0.02** |
| **Male weight** | 0.14 | 0.16 | 0.93 | 0.32 |
| **Male age** | 0.00 | 0.10 | 0.04 | 0.96 |

**Further details of the results on genotype-based and lab-reared vs-wild caught differences in male antennal morphology (referred to the Results section in the main text)**

In wild-caught males, male color morph did not affect antennal length (F_(1, 137)_ = 0.27, *P* = 0.61), area (F_(1, 137)_ = 0.03, *P* = 0.87), or the number of lamellae (F_(1, 137)_ = 1.87, *P* = 0.17). For laboratory-reared males, the color genotype affected the area of antenna (F_(2, 99)_ = 5.78, *P* = 0.004) but not antenna length (F_(2, 99)_ = 0.32, *P* = 0.73), or the number of lamellae (F_(2, 99)_ = 1.56, *P* = 0.22). WW antennae were wider compared to Wy (lm; estimate = -0.15 ± 0.05, t = -2.81, *P* = 0.006) and yy (lm; estimate = -0.19 ± 0.05, t = -3.49, *P* = 0.001) males. While male weight did not affect the number of lamellae (F_(1, 99)_ = 0.38, *P* = 0.54), the heavier the male, the longer (lm; estimate = 0.18 ± 0.04, t = 4.63, *P* < 0.001) and wider (lm; estimate = 0.17 ± 0.03, t = 6.52, *P* < 0.001) the antenna. Regardless of the origin of the male, antennae had an equal number of lamellae (F_(1, 260)_ = 1.92, *P* = 0.17) and no significant interaction of male origin with color morph was found for antenna length (F_(1, 260)_ = 0.10, *P* = 0.75), area (F_(1, 260)_ = 0.90, *P* = 0.34), or lamellae (F_(1, 260)_ = 0.73, *P* = 0.39), excluding the morph-specific laboratory effect. Wild-caught males were characterized by longer (lm; estimate = 0.14 ± 0.05, t = 3.04, *P* = 0.003) and wider (lm; estimate = 0.08 ± 0.03, t = 2.46, *P* = 0.01) antennae compared to laboratory-reared males.

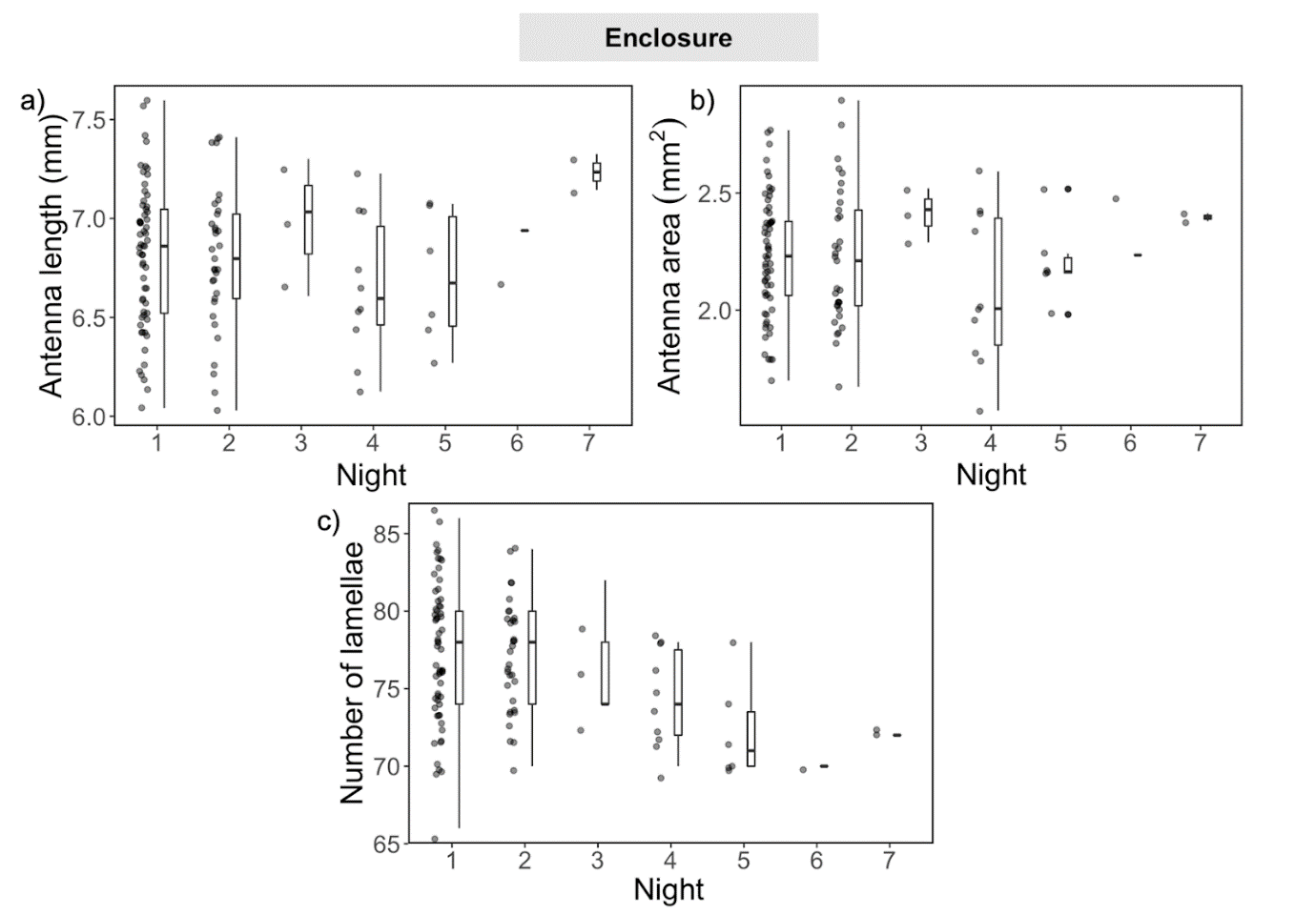

FIGURE S3	In the enclosure, a) the length and b) area of the male antennae did not affect their order of arrival to female traps whereas c) males with higher number of lamellae were more likely to be attracted on the first night, rather than subsequent nights. For each night of the experiment, the antenna trait is described by the individual measurement (i.e., the dots) and as summary of the male population for the night (i.e., the boxplot). The top and the bottom of the box represent the first and third quartiles, respectively. The line within each box indicates the median and the vertical lines extending from the box (whiskers) represent the maximum and minimum values.
